# Thermal laser ablation with tunable lesion size reveals multiple origins of seizure-like convulsions in *Caenorhabditis elegans*

**DOI:** 10.1101/2021.01.27.428539

**Authors:** Anthony D. Fouad, Alice Liu, Angelica Du, Priya D. Bhirgoo, Christopher Fang-Yen

## Abstract

Laser microsurgery has long been an important means of assessing the functions of specific cells and tissues. Most laser ablation systems use short, highly focused laser pulses to create plasma-mediated lesions with dimensions on the order of the wavelength of light. While the small size of the lesion enables ablation with high spatial resolution, it also makes it difficult to ablate larger structures. We developed an infrared laser ablation system capable of thermally lesioning tissues with spot sizes tunable by the duration and amplitude of laser pulses. We used our laser system in the roundworm *C. elegans* to kill single neurons and to sever the dorsal and ventral nerve cords, structures that are difficult to lesion using a plasma-based ablation system. We used these ablations to investigate the source of convulsions in a gain-of-function mutant for the acetylcholine receptor ACR-2. Severing the ventral nerve cord caused convulsions to occur independently anterior and posterior to the lesion, suggesting that convulsions can arise independently from distinct subsets of the motor circuit.

## Introduction

Laser ablation is a powerful tool for creating specific cellular or tissue lesions in various model organisms^1-3^. In the roundworm *C. elegans*, laser killing has been used to study cell function^4-8^, cell-cell interactions^9-12^, and the effects of severing nerve fibers^13-15^.

Ablations in *C. elegans* are normally performed using localized plasma generated by pulsed lasers with high peak power, coupled through a microscope objective^1^. Since plasma generation occurs only at the laser focus, plasma-mediated ablation has high spatial resolution, on the order of the wavelength of light used.

In our recent studies of the *C. elegans* locomotor circuit^16^ we sought to sever the ventral nerve cord (VNC), which has a diameter ranging from 2 to 4 microns in the adult^17^. We initially sought to use a 3-5 nanosecond duration, nitrogen-laser-pumped dye laser (MicroPoint, Andor Technologies) for this microsurgery. However, we encountered three problems. First, ablation of deep structures in the adult worm did not work reliably, possibly due to depth-induced optical aberrations. Second, since the VNC is much larger than the lesion size, many laser pulses were required, at varying positions along the lesion site, in order for the VNC to be completely severed; this procedure was difficult and time-consuming. Third, since the VNC is located just beneath the worm’s cuticle (skin), inadvertent lesion of the cuticle often caused the animal to burst due to release of the worm’s internal hydrostatic pressure^18^. For these reasons, we found severing the VNC using a nanosecond dye laser system to be difficult and impractical.

An alternative to plasma-mediated laser ablation methods is to induce a local elevation of tissue temperature through absorption of a focused infrared laser beam. Thermally mediated laser ablation has a long history in biology and biomedicine^19^; recently it has been used to kill *C. elegans* embryonic cells^20^ and cauterize tissue in *Drosophila*^21^.

Here we describe and characterize an infrared laser method for performing thermal ablation in *C. elegans*. Compared with plasma-mediated ablation, our thermal ablation method has the advantage of a tunable lesion size through changes in the laser pulse duration and/or power, and can be used to easily lesion relatively large structures such as the ventral nerve cord without puncturing the cuticle. In addition, this method can create lesions throughout an adult animal, possibly because it is relatively insensitive to sample-induced optical aberrations.

We used our infrared laser ablation system to probe the generation of aberrant motor excitations in worms with a gain-of-function mutation in the acetylcholine receptor ACR-2. This mutation is analogous to a mutation responsible for congenital myasthenic syndrome in humans and results in seizure-like convulsions in worms due to hyperactivation of the B-type motor neurons^22,23^. It is unclear whether these convulsions arise from one or multiple sources. We found that lesioning the VNC caused convulsions to arise independently in the animal anterior and posterior to the lesion. These results imply that convulsions do not arise from a single locus but can originate in multiple portions of the circuit.

## Results

### The thermal lesion increases in size with laser pulse duration

To induce thermal lesions in *C. elegans*, we modified a pulsed infrared laser system (see Methods) previously used to induce heat shock in single cells ^24,25^ (Fig. 1). First, we sought to test the ability of this laser to ablate specific neurons without damaging neighboring cells. We also aimed to determine the approximate lesion size as the pulse duration and/or number of pulses was increased.

**Fig. 1:**
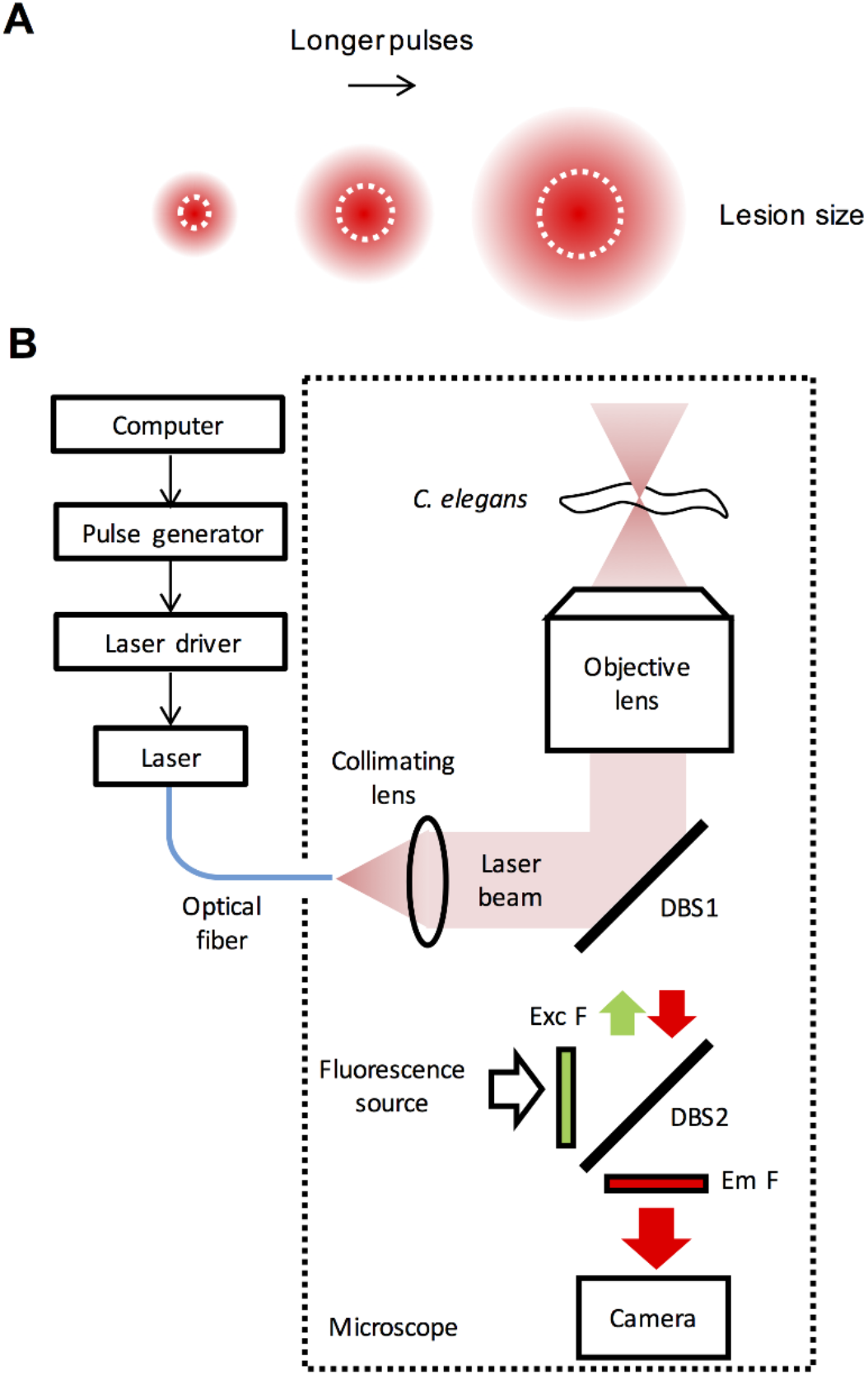
Thermal ablation system. (A) Basic principle of thermal ablation with variable lesion size. Increased duration and/or number of pulses increases the total amount of heat delivered, thereby increasing the spatial extent of thermal damage. (B) Infrared laser ablation system. A computer and pulse generator trigger the infrared laser to produce pulses of tunable length. A microscope objective focuses the laser beam within the worm, producing localized thermal damage.

We performed thermal ablations in a worm in which the red fluorescent protein wCherry is expressed in a subset of ventral cord motor neurons via the transgene P*acr-2::wCherry*. We targeted the VB9 motor neuron with a single laser pulse of varying duration and evaluated the fraction of worms in which VB9 and/or the neighboring motor neurons VA8 and DB6 remained alive 24 h later (Fig. 2).

**Fig. 2:**
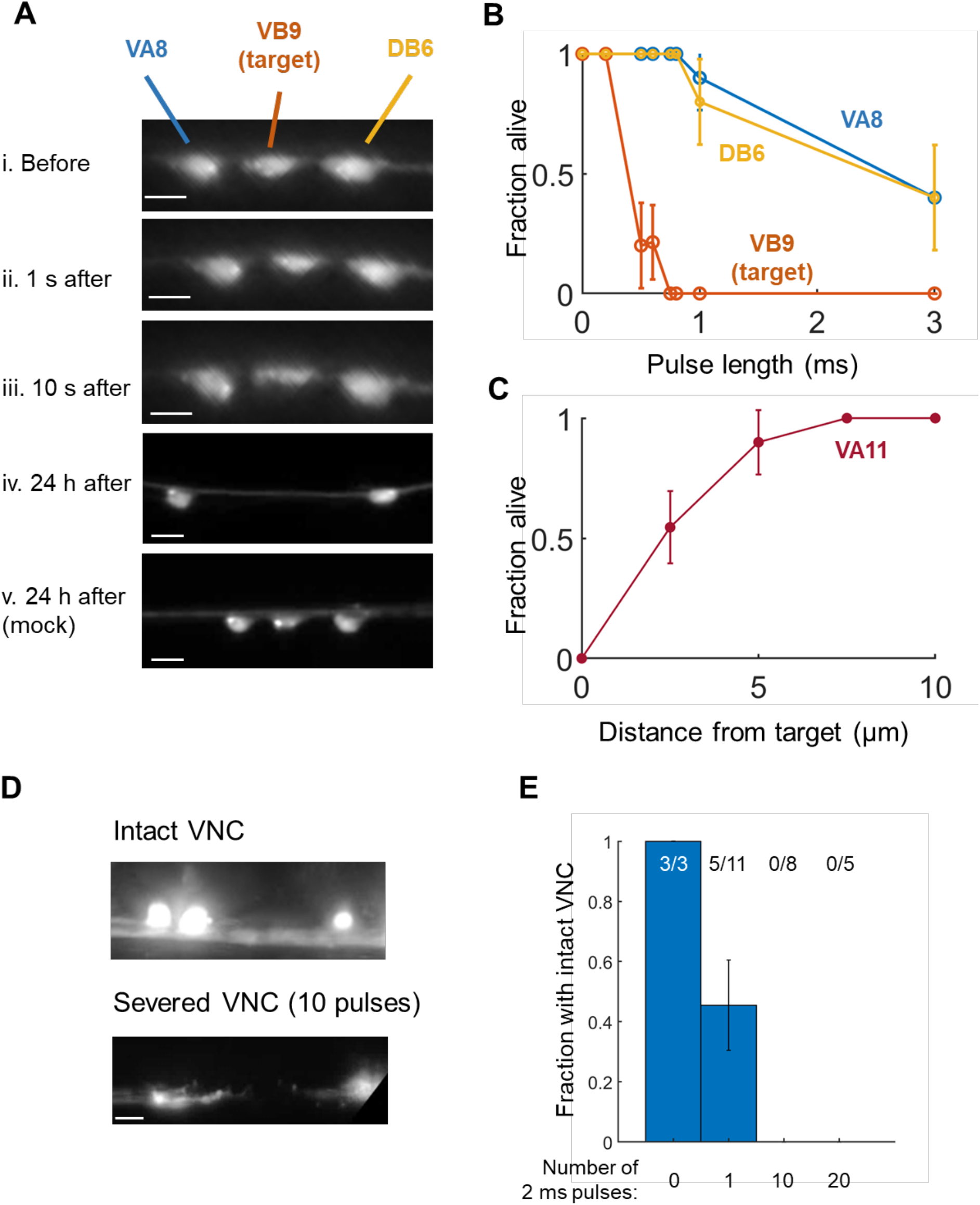
Lesion of specific cell bodies and nerve bundles using infrared laser surgery. (A) Three VNC motor neurons (VA8, VB9, DB6) immediately prior to irradiation of VB9 with a single 0.6 ms pulse (i), 1 s after irradiation (ii), 10 s after irradiation (iii), and 24 hours after irradiation (iv). All scale bars are 5 μm. VA8, VB9, DB6 shown in a mock ablated animal (v). Both animals are day 1 adults. Brightness has been increased 40% in both iv and v. (B) Fraction of worms in which the indicated neuron was alive 24 hours after irradiation of VB9 with a single pulse of the specified length. N = 5-8 worms per condition, except for pulse lengths of 0 ms, for which N = 2 worms. (C) Fraction of worms in which VA11 was alive 24 hours after irradiation of the VNC at the specified distance away from the center of the VA11 cell body with a single 0.8 ms pulse. N = 5-6 worms per condition, except for 2.5 µm which had N=11 worms. (D) Red fluorescence image of the ventral nerve cord shown after 0 or 10 pulses of length 2 ms were applied and the worm was allowed to recover for at least four hours. The VNC was severed after application of a train of 10 pulses. Both images are shown with brightness increased by 40%. (E) Fraction of worms in which the VNC was not completely severed after application of 0, 1, 10, or 20 pulses. Error bars are the standard error of the sample proportion.

Immediately after a single 0.75 or 0.8 ms pulse, the targeted VNC motor neuron appeared to shrink and its fluorescence faded, while the morphology of neighboring cell bodies appeared unchanged (Fig. 2A). When we assayed 24 h later, we found that in every trial, VB9 was killed while the two neighboring neurons remained intact. Longer pulse durations increased the probability of killing VA8 and DB6 in addition to the targeted neuron, while shorter pulse durations failed to reliably kill VB9 (Fig. 2B). These results suggest that in our system, a pulse duration of 0.75 - 0.8 ms is well suited to killing an individual VNC neuron.

To quantify the approximate cell killing radius of a single 0.8 ms laser pulse, we conducted pulsed illuminations with the focus at distances ranging from 0 to 10 μm from the center of the wCherry-labeled VA11 cell soma. The neuron VA11 was almost never killed when the pulse was centered at least 5 μm away from the center of the cell body, and was killed in about half of animals when the pulse was applied 2.5 μm from the center of the cell body (Fig. 2C), corresponding approximately to the edge of the cell body.

Next, we tested the system’s ability to lesion the VNC of the adult worm (Fig. 3A). Although laser dissection is routinely performed on *C. elegans* neuronal processes^1^, to our knowledge it had not previously been reported other than in our recent publication^16^ on the VNC, a bundle with a total width ranging from 2 to 4 µm^17^. We performed laser surgeries on the VNC in a worm expressing GFP in all cholinergic neurons (see Methods) and assessed lesions by imaging fluorescence 4 h after surgery. We found that a single 2 ms pulse, which is sufficient to kill a neuron cell body (Fig. 2B), did not reliably cut the nerve cord (Fig. 2E). However, trains of 10 or 20 pulses, each of length 2 ms, cut the nerve cords in all worms tested (Fig. 2E).

**Fig. 3:**
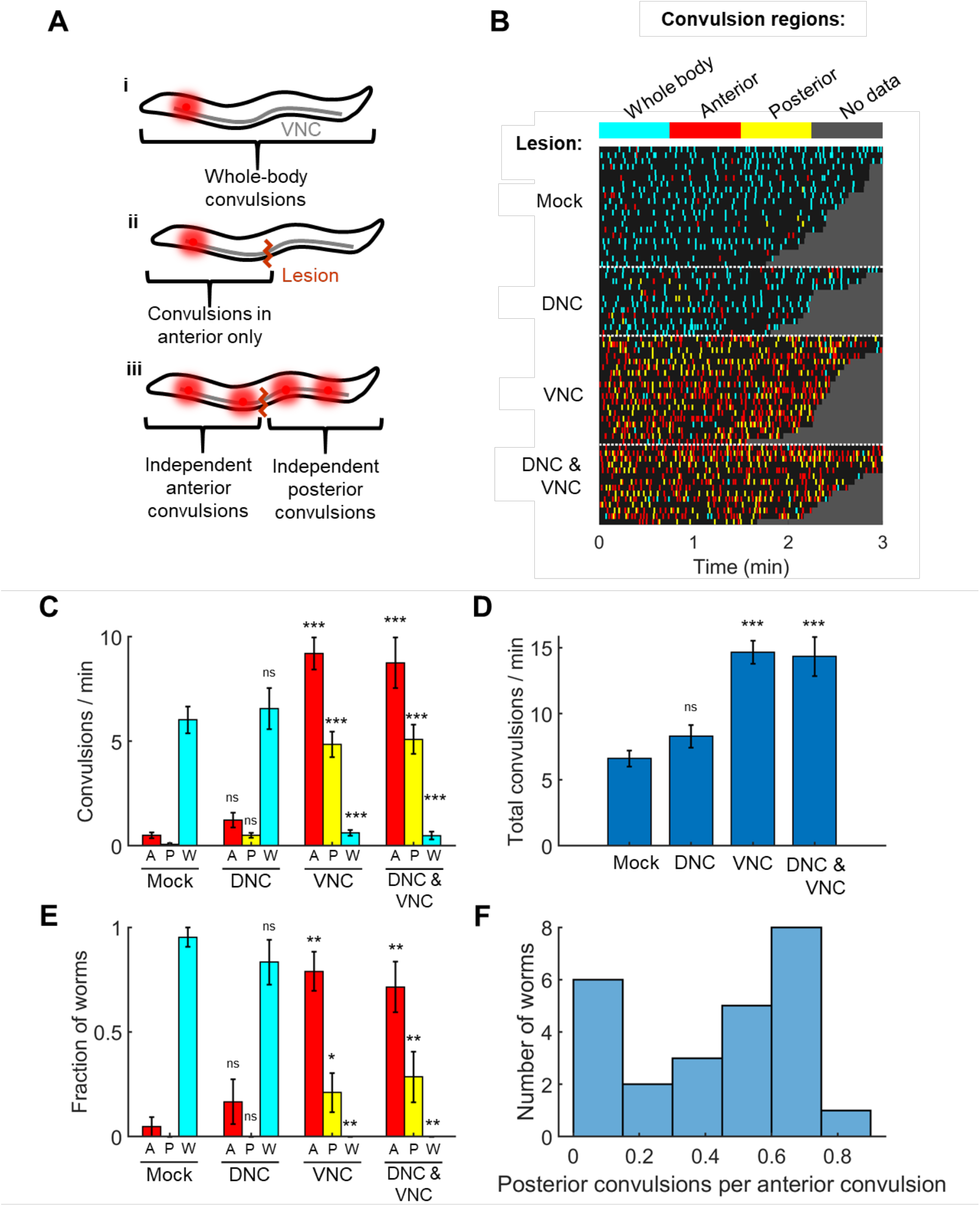
Lesioning the ventral nerve cord of *acr-2(gf)* mutants affects patterns of convulsions in the animal’s anterior and posterior. (A) Model in which convulsions originate from a single locus (red spot) and then propagate throughout the VNC (i), causing a whole-body convulsion. (ii). Severing the VNC should result in convulsions on the side of the lesion containing the locus. (iii) Model in which convulsions can arise from multiple loci. If at least one locus is located anterior to the lesion and at least one locus is located posterior to the lesion, severing the VNC may cause the animal’s anterior and posterior to convulse independently. (B) Timing and classification of convulsions in worms with no lesion (N=21), mid-body lesions to the DNC only (N=12), VNC only (N=19), or VNC and DNC (N=14 worms). Convulsions were classified as occurring in the whole body (cyan), anterior only (red), or posterior only (yellow). Each row corresponds to one worm. Grey region follows the end of each recording. (C) Mean and SEM rates of anterior (A, red), posterior (P, yellow), and Whole-body (W, cyan) convulsions with each lesion type. * p<0.05, ** p < 0.01, *** p<0.001, one-way ANOVA with Bonferroni-corrected post-hoc comparisons to the mock control mean of the same color. (D) Total convulsion rate (A + P + W) for the worms in each lesion group. *** p < 0.001, one-way ANOVA with Bonferroni-corrected post-hoc comparisons to the mock control. (E) Fraction of worms exhibiting primarily anterior (A, red), primarily posterior (P, yellow), or primarily whole-body (W, cyan) convulsions for each nerve cord lesion. *p<0.05, ** p < 0.01, two-proportion Z test against the mock control bar of the same color. (F) Histogram showing the rate of posterior convulsions (shown as a fraction of anterior convulsions) among the 25 worms with VNC or DNC & VNC lesions that were classified as having primarily anterior convulsions; 1 out of the 25 worms did not have any posterior convulsions (included in the leftmost bar).

These results show that pulsed infrared laser-based ablation can be used to kill specific *C. elegans* cells, that the lesion size increases with pulse duration, and multiple pulses can be used to lesion larger structures such as the VNC.

### Convulsions in *acr-2 gain-of-function* mutants do not arise from a single locus

We used our infrared ablation system to probe the nature of aberrant motor excitations in *C. elegans acr-2 gain-of-function (gf)* mutants. This mutation causes worms to exhibit sporadic whole-body muscle contractions (Video S1). These behaviors have been described as epileptic-like convulsions and arise from hyperactivity in B-type motor neurons^22,23^.

In some cases of epilepsy in humans, seizures originate from a localized region (focus) and then propagate to a larger region in the brain; lesion of the area important to the genesis of seizures can lead to cessation of seizures^26,27^. In other cases of epilepsy, seizures arise from multiple regions ^28^.

In *C. elegans acr-2(gf)* mutants it is not known whether the aberrant excitations originate at a specific locus in the nervous system (e.g. a particular neuron or sub-circuit), or from multiple loci, or from throughout the B type motor network (Fig. 3A). If convulsions originate from a focus, severing the VNC should result in convulsions being observed either anterior or posterior to the lesion (Fig. 3A(ii)), depending on the location of the focus, but not both anterior and posterior. If instead convulsions originate from multiple foci, with at least one anterior of the lesion and at least one posterior to the lesion, or from the motor network as a whole (Fig. 3A(iii)), we would expect severing the VNC in the midbody to result in independent convulsions anterior and posterior to the lesion.

To address this question we lesioned the VNC and/or dorsal nerve cord (DNC) near the middle of the body in worms with the *acr-2(gf)* mutation. Worms were then recovered to NGM plates. Four hours after surgery, we recorded the convulsion behavior of each worm. Preliminary observations indicated that after ablation, convulsions persisted anterior and posterior to the lesion site; therefore we scored each convulsion as occurring in the whole body, anterior half of the body only, or posterior half of the body only (Fig. 3B).

Worms in the mock ablation group, and in worms in which only the DNC was severed, predominantly exhibited whole body convulsions (Fig. 3B-C) and a small number of anterior convulsions. By contrast, worms in which the VNC only or both the VNC and the DNC were severed showed a mixture of anterior, posterior, and whole-body convulsions, with anterior convulsions the most common. The total rates of convulsions for worms with the VNC or both nerve cords lesioned were significantly higher than in mock controls (Fig. 3D).

We observed considerable inter-worm variability in the propensity for different types of convulsions (Fig. 3B-C). To explore the patterns of convulsions on an individual animal basis, we classified each worm as having primarily anterior, posterior, or whole-body convulsions according to the most common convulsion type for that animal. A large majority of the worms in the mock ablation (control) group, and of the worms in which only the DNC was severed, exhibited primarily whole-body convulsions (Fig. 3E). By contrast, most of the worms in which the VNC or both the VNC and the DNC were cut showed primarily anterior convulsions, and very few primarily posterior convulsions (Fig. 3E).

Worms in which convulsions occurred primarily in the anterior after mid-body lesions may either have only anterior foci or may have additional posterior foci that are less active than those in the anterior. We examined the occurrence of posterior convulsions in worms with either the VNC or both nerve cords lesioned that were classified as having primarily anterior convulsions. Only 1 of the 25 worms in this group showed no posterior convulsions at all. The remaining animals exhibited posterior convulsions at variable rates relative to the anterior convulsions (Fig. 3F).

These results suggest that in most *acr-2(gf)* worms, seizure-like activity does not originate from a single focus, but instead may arise from either the anterior or posterior of the animal. However, our finding that posterior-only convulsions are less frequent than anterior-only convulsions suggests that convulsions are generated more actively in the anterior half of the animal. Finally, the finding that cutting the VNC led to anterior-only and posterior-only convulsions, while cutting the DNC did not, suggests that the abnormal excitations in *acr-2(gf)* mutants propagate primarily via the VNC.

## Discussion

We have presented thermal lesioning with an infrared laser as an alternative to conventional microsurgery in *C. elegans*. Advantages of our system relative to plasma-mediated laser ablation include the ability to generate continuously variable lesion sizes and the relative insensitivity to optical aberrations, which makes it possible to perform lesions throughout the adult. The thermal nature of the lesions may allow for the retainment of more structural integrity in ablated tissue as proteins denature rather than vaporize as in plasma-mediated ablation. This may help to avoid problems with the worm rupturing due to laser damage to its hydrostatically pressurized cuticle.

One disadvantage of our system is that the smallest lesion size is larger than that of plasma-mediated ablation. In addition, it is possible that neighboring cells and tissues may experience heat shock or other temperature-related damage due to heat diffusing from the thermal lesion.

We used our thermal lesioning system to examine the origins of convulsions in *acr-2(gf)* mutants and showed that convulsions typically do not originate from a single locus. It has been shown that *acr-2(gf)* mutants have hyperactive B-type motor neurons^22,23^.

How does general hyperactivation lead to sporadic seizures that can originate at multiple foci? In humans, neuronal hyperactivity can drive neurons towards a seizure threshold ^29^; neurons and neuronal circuits closest to the threshold may be most susceptible to initiating seizures. It is possible that *C. elegans acr-2(gf)* convulsions also occur when excitation within a focal structure exceeds a threshold. Because neurons may function normally except when excitation exceeds a certain threshold, persistent hyperactivity leads to essentially normal behavior most of the time, with occasional convulsions. This model is consistent with the observations that stimulation of B motor neurons optogenetically triggers a convulsion (Fouad, Liu, and Fang-Yen, unpublished data), as does activation of the premotor interneurons AVB^23^, which are coupled to the B type motor neurons^30^.

We observed that lesioning the VNC or the VNC and DNC together increased the total rate of convulsions. One possible explanation for this result is that the laser-induced damage to VNC motor neurons caused calcium influx, as has been found for other *C. elegans* neurons after injury^31-33^. The resulting elevated excitability in these cells may have increased the frequency of convulsions.

One outstanding question is the extent to which severed nerves can regain function after thermal lesioning, as they have been observed to do after plasma-mediated laser surgery^15,34^. We limited the time between surgery and behavioral experiments to 4 hours because we observed possible regrowth of the nerve cord in some animals left overnight (see Methods). It remains unclear whether the dozens of nerve processes in the VNC are capable of correctly repairing their connections after lesions, and how the nature of damage delivered by various ablation systems affects the subsequent recovery.

## Methods

### Thermal ablation system

The laser system (Fig. 1A) is similar to that previously used for local heat shock^24,25^, and is built around an inverted microscope (Nikon TE-2000). A thermoelectrically cooled, fiber-coupled diode laser (Fitel FOL1425RUZ-317, 1480 nm wavelength) generates a beam that emerges from the fiber end and is collimated by a lens. The beam reflects from a dichroic beam splitter, enters an objective lens (Nikon PLAN APO 60X, NA 1.4, oil immersion), and is focused onto a sample slide (Fig. 1B). A computer, pulse generator, and laser current source (Opto Power OPC-PS03-A) drive the laser to generate pulses of the specified frequency and duration.

We made several modifications to the previous heat shock system^24,25^ to adapt it for laser ablation. First, we replaced the original collimating lens (focal length f = 75 mm) with one of smaller focal length (f = 11 mm) in order to create a smaller diameter beam, allowing a larger fraction of the beam power to transmit through the objective and reach the sample. Second, the laser power output was increased from 260 mW to 400 mW. Third, we adjusted the pulse duration and count as described above.

### Strains

*C. elegans* were maintained using standard methods^35^. The following strains were used in this study: YX148 (qhIs4[P*acr-2::wCherry*]; qhIs1[P*myo-3::NpHR::CFP*]) (for cell ablation studies), YX200 (qhIs4[P*acr-2::wCherry*]; qhIs1[P*myo-3::NpHR::CFP*]; *acr-2(n2420))* (for convulsion studies), YX173 (qhIs4[P*acr-2::wCherry*]; qhIs1[P*myo-3::NpHR*]; vsIs48[P*unc-17::*GFP]) worms (for nerve cord ablation studies). Although some strains expressed the inhibitory opsin NpHR^36^, it was not functional as our plates did not contain the cofactor all-*trans* retinal.

### Laser surgery procedures

For nerve cord lesion experiments, day 1 adult YX200 or YX173 worms were immobilized on agar pads using polystyrene beads^37^ and mounted on the microscope. The wCherry or GFP-labeled nerve cord was imaged under RFP or GFP optics and illuminated with a train of up to 20 laser pulses with 2 ms duration. Worms were transferred to fresh seeded plates and allowed to recover for 4 hours before behavioral and fluorescence imaging, as we observed that severed nerve cords sometimes appeared to regrow when left overnight (not shown). Mock controls were mounted on the system but not illuminated with the laser.

For cell killing experiments, fourth larval stage YX148 worms mounted as described above, subjected to one pulse of infrared illumination at the targeted site, and then recovered to seeded plates and allowed to recover and grow overnight. Animals were day 1 adults during behavioral assays the following day.

### Behavioral and imaging assays

After ablations and recovery, YX200 worms were picked to a new unseeded plate for 5 minutes before the start of recording. Unseeded plates were used to improve the visibility of worms for behavioral scoring. The unseeded plates were imaged under RFP fluorescence optics on a stereo fluorescence microscope (Leica M165FC). We briefly recorded images of the worms at approximately 60X magnification to confirm that the targeted nerve cords were lesioned. We did not proceed with further analysis of any animal in which the targeted nerve cords did not appear to be completely severed.

We acquired image sequences of the fluorescent motor neurons at approximately 10X magnification at 10 frames per second for 2-3 minutes per worm using a CMOS camera (Imaging Source DMK 23GP031). Fluorescence imaging aided in scoring because the visualization of fluorescent neurons made it easier to assess local deformations during convulsions (Videos S1, S2).

To score convulsions, we used a simple application written in C++ to play video recordings in random order and collect input from the scorer. The scorer marked the time of each convulsion and identified it as occurring in the animal’s anterior, posterior, or whole body. All scoring of convulsions was carried out by viewers blinded to the experimental group.

Imaging and scoring of neuronal death in YX148 worms or nerve cord lesions in YX173 worms was carried out as previously described^16^. Briefly, worms at the indicated timepoint were imaged for red or green fluorescence on a compound microscope (Leica DMI6000B), and neurons were identified using their stereotypic positions and other features.

## Acknowledgements

We thank Mei Zhen, Yishi Jin, and Alexander Gottschalk for providing strains. Some strains were provided by the by the CGC, which is funded by NIH Office of Research Infrastructure Programs (P40 OD010440). CFY and ADF were supported by the National Institutes of Health (R01 NS-084835). AL was supported by the Vagelos Scholars Program in the Molecular Life Sciences.

## Supporting Information

**Video S1: Convulsions in a YX200 *acr-2(gf)* worm with no lesions**. Video acquired using RFP fluorescence optics and shown at 1.5 times the original frame rate. Field of view is approximately 1 mm x 1.5 mm. Whole body convulsion indicated with ‘W’.

**Video S2**: **A YX200 worm with mid-body lesions to both the VNC and DNC exhibiting independent anterior and posterior convulsions**. Anterior of the worm is to the upper left. Video acquired using RFP fluorescence optics and shown at 1.5 times the original frame rate. Field of view is approximately 1 mm x 1.5 mm. Annotations denote convulsions scored as occurring in the anterior (‘A’) or posterior (‘P’).

